# Mechanistic Model of Telomere Length Homeostasis in Yeast

**DOI:** 10.1101/2021.10.02.462846

**Authors:** Ghanendra Singh, K. Sriram

**Affiliations:** Center for Computational Biology, Indraprastha Institute of Information Technology Delhi, 110020, India

**Keywords:** Telomere Length (TL), Telomerase (Telo), Rif1, Tel1, Cdc13

## Abstract

Cells maintain homeostatic telomere length, and this homeostatic disruption leads to various types of diseases. Presently, it is not clear how telomeres achieve homeostasis. One of the prevailing hypotheses is a protein-counting model with a built-in sensor mechanism that counts proteins that directly regulate the telomeric length. However, it does not explain telomere length regulation at the mechanistic level. Here, we present a mathematical model based on the underlying molecular mechanisms of length regulation needed to establish telomere length homeostasis in yeast. We perform both deterministic and stochastic simulations to validate the models with the experimental data of Teixeira et al., rate-balance plot, and phase plane analysis to understand the nature of dynamics exhibited by the models. For global analysis, we constructed bifurcation diagrams. The model explains the role of negative and positive feedback loops and a delay between telomerase and telomere-bound proteins, leading to oscillations in telomere length. We map these in-silico results to Teixeira’s proposition of telomeres making a transition between extendible and non-extendible states.

## Introduction

During DNA replication, DNA polymerases can synthesize new DNA strands only in 5’ to 3’ direction. It requires a primer that provides a free 3’ OH group at the end but the replication machinery fails to duplicate the 3’ ends of the chromosomes due to the non-availability of the OH group which leads to the shortening of chromosomes during each cell division for linear chromosomes and this problem does not exist with circular chromosomes. This property of DNA polymerase creates the end replication problem [45, 44] in eukaryotes. Eukaryotic cells have developed a novel solution involving specialized structures called telomeres at the end of their chromosomes [4] to deal with incomplete replication compared to prokaryotic organisms. Telomeres are repetitive sequences present at the end of the chromosomes protecting them from fusion and recombination [36, 64]. Telomeric DNA consists of a G-rich strand (50-30) and a C-rich strand (30-50) and is made up of short, simple repeats [65]. Telomere length shortens during cell division but an enzyme known as telomerase maintains the telomere length by elongating it throughout many cell divisions to keep it at equilibrium [21]. Telomeres shorten continuously in human somatic cells because of the end replication problem but telomere length is maintained in tumor cells, germline cells, and unicellular organisms like yeast by a mechanism expressing telomerase enzyme which elongated the 3’ ends of telomeres [11, 21, 29, 60]. This telomere length maintenance is necessary in the germline but may not be in somatic cells for survival. Lower eukaryotes have short telomeres whereas telomeres of vertebrates contain longer DNA (5 to 15kb) with large size distributions [22]. So, it is reasonable to assume that mechanisms exist in these organisms to maintain telomere length at some steady state for cell survival.

Biological systems can self-regulate their internal state to maintain stability and function optimally despite external fluctuations. This process is known as homeostasis in contrast to adaptation in which organisms can adapt themselves to an externally changing environment. Many homeostatic processes require negative feedback regulation to maintain a specific parameter around a particular range. Cells have to maintain their telomere length within a range for healthy proliferation. How does the telomere length is maintained?.

A protein counting model was proposed earlier by [39] for budding yeast *S. cerevisiae* that by counting the number of telomeric binding protein Rap1p length of the telomeric DNA can be measured. In this model, a higher number of telomere bound proteins negatively regulate telomerase activity on that telomere. Telomere length homeostasis is achieved through a dynamic switching of telomeres between two uncapping capping cycle states as mentioned earlier by [5]. Later another probabilistic model of switching between short telomeres (extendible) and long telomeres (non-extendible states) with a higher probability of switching of short telomeres was proposed by [55]. This regulatory mechanism of telomere length homeostasis was described as a balance between elongation and shortening state of telomeres. How does this transition in the state take place at the molecular level? How this telomere length distribution is generated? What are the factors that determines whether a telomere will be shortened or elongated during S phase and how does this leads to equilibrium of telomere length in yeast?

Recently an alternative model the replication fork model was hypothesized by [20] which explains role of replication fork progression and origin firing in regulating telomere length. The existing protein counting model [39] is able to explain the negative inhibitory effect of telomere binding proteins on telomere length based on experimental data but still based on the recent experimental findings and proposed molecular mechanisms, it does not account for origin firing and role of DNA replication in telomere length homeostasis. It does not explain the mechanisms regulating telomere length at the molecular level. It does not explain the role of DNA replication particularly of replication fork in telomere length.

In *S. cerevisiae*, telomeres is bound by ssDNA binding protein Cdc13 [33, 43] and Stn1 and Ten1 proteins [19, 18]. Rap1p counting is Rif protein counting as a shortening of telomere is proportional to the bound Rif1/Rif2 proteins as shown earlier as counting of Rap1 protein [31]. Rap1p protein binds to double-strand telomere repeats [52]. Telomeric proteins protect telomeres and regulate telomerase access to the telomere [47, 58]. How they carry out these functions is critical to understanding length regulation. Similarly, in mammals, we have Trf1, Trf2, Pot1, and Tin2 proteins and their knockdown leads to excessive telomere elongation [57, 34, 63, 53]. Only 20 number of telomerase are present in yeasy whereas human cancer cells consists of around 250 number of telomerase in each cell [42, 61]. Rif1 negatively regulate telomere length in *S. cerevisiae* [24]. Interestingly, when a telomere is artificially shortened, the telomere-proximal origin fires more efficiently [3]. Cdc13 and Est1 recruit telomerase to the telomere. Cdc13 binds telomerase through interaction with Est1 [14]. In S. *cerevisiae* Rap1p protein binds not only to the telomeric region but also to recognition sites within chromosomes [17].

From a biological circuit perspective which requires three things: sensor, transducer, and effector. Proteins or elements which have to be counted act as sensor or detector, proteins or complexes that send this information act as a transducer and finally the proteins or complexes that take action to elongate or shorten the telomeres. Studies from [3] described in detail the molecular interactions of telomeric proteins. The underlying molecular mechanism through which the signal is relayed from any sensor proteins like (Rap1, Rif1, and Rif2) to the transducer proteins (MRX and Tel1) to establishes telomere length homeostasis [35]. Molecular interactions in the budding yeast protein counting machinery were described by [27]. Telomere is extended by telomerase in the late S phase [38]. Rif1 negatively regulates telomeres in the *S. cerevisiae*. Rif1 recruits PP1 to inhibit the recruitment of Tel1 kinase which suppresses the lengthening mechanism of telomerase [28].

Earlier mathematical models that were proposed have dealt specifically with a particular problem statement. A probabilistic model based on binomial distribution for telomere end replication problem was given by [32] which predicted the number of deletion a telomere undergoes before replicative senescence. It described the characteristics of the passive incomplete replication model for telomere loss assuming no elongation by telomerase. Another theoretical model was proposed for telomere length regulation given by [30] which proposed a mechanism to maintain telomeres at a constant length, how mutations lead to elongation of telomeres, introduced inhibition of telomerase which causes telomeric shortening, and explained experiment results of addition of oligonucleotides to increase telomere length. Later a stochastic model of telomere dynamics built on the protein counting model describing the role of short telomere in triggering senescence and showed that length of the short telomere is a key genetic marker determining senescence onset in *S. cerevisiae* [62]. Other models for the variability of senescence onset [54, 48, 49, 46, 2]. One of the studies built on the t-loop structure as the molecular basis of its model, a structure observed in eukaryotic telomeres [23, 51] but not in yeast. But the above models so far did not discuss the mechanisms that regulate telomere length at the molecular level. Studying molecular mechanisms will help us to relate telomere regulation with the dynamics of DNA replication and understand the role of DNA replication checkpoint kinases. How cells actually count telomere-bound proteins? How these processes regulate each other? Next to predict what might be the molecular dynamics operating in the cancer cells or stem cells regulate telomere length in comparison to somatic cells regulation.

Here we present a mathematical model based on molecular mechanisms of telomere length regulation. We describe how the telomere length might be oscillating between telomere elongation and shortening state and compare with earlier model of switching between extendible and non-extendible states [55] and we also explain how telomere length distribution must be getting generated and maintained.

In the below Methods section, we have discussed a step-wise process of designing the biological circuit in detail of telomere length homeostasis starting from a simple 1D model of Telomerase and telomere bound protein Rif1 showing how the presence of low telomerase concentration in somatic cells or high telomerase concentration in stem cells can be easily modeled and can predict how injecting externally some telomerase can lead to elongation of telomere length. Next in the 2D model, we discuss the role of negative feedback of Rif1 on Telomerase giving rise to telomere length homeostasis. Later in the 3D model inclusion of DNA checkpoint kinase results in damped oscillation and finally in the 4D model addition of telomerase recruitment factors gives rise to sustained oscillations of telomere length.

## Materials and Methods

Python is used to write all the source code related to mathematical models. In python scipy.integrate.odeint module is used for integrating ODEs. It uses a solver LSODA (solver for ODE) algorithm which adapts the step size. This means that it estimates the integration error, e.g., by comparing the result of two simple integrators (which are intelligently intertwined to save runtime). This estimate is then used to adapt the step size such that the estimated error is below a user-defined threshold specified by the arguments rtol and atol. These arguments allow you to control the precision. Xppaut [13] is used for stochastic simulation and bifurcation analysis.

### Telomere length equation

An Interaction between DNA replication machinery, DNA damage response factors, and telomeric proteins that recruits and regulates telomerase indicates how does protein counting mechanism works [52]. Based on the experimental data by [17, 50], Rap1 binds to telomeric dsDNA, potentially at a density of 1 in every 18 bp [59] the telomere length is maintained at 250 to 350 base pairs. Given these parameters, it’s fair to consider that a normal telomere can have 14–20 Rap1 binding sites and Rap1 proteins on which many no. of Rif1 and Rif2 bound. So we have provided a simple expression to determine the telomere length based on the protein counting mechanisms by counting the no. of Rif1 proteins bound to the telomere. Telomere Length (TL) = No. of base pairs each Rap1 binds to dsDNA (18bp) multiplied by the no. of Rap1 proteins (nR) which binds to dsDNA.

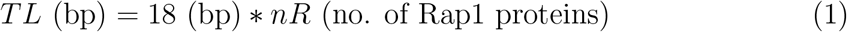

Using equation 1, we have calculated the telomere length distribution that relates with the experimental results [17] and previous Teixeira’s (TL) distribution [55].

## Results

### 1D Futile Cycle Model

We first construct a simple 1D differential equation model to get the telomere length maintained at the steady state level in the normal condition. In S. *cerevisiae* telomeres are maintained at an equilibrium length of 300 base pairs (bps) [55]. The steady state telomere length (TL) throughout this paper for S. *cerevisiae* is taken between 250-350 (bps). One of the well studied protein involved in telomere length dynamics is the telomere bound protein [Rif1] modulated by the reverse transcriptase enzyme telomerase. We have taken telomerase as a parameter that modulates the telomere length during DNA replication. This model has no regulation and captures only the production and loss of [Rif1] protein by telomerase and therefore it is a model of “futile” cycle, The model is given below by the following equation.

Rap1 protein binds to the double strand telomeric DNA. Rif1/Rif2 binds together on the Rap1 protein. Rif1 acts as a sensor to transmit the signal downstream to kinase proteins. Rif1 has a key role in DNA replication, regulation and many other import processes as discussed by [26, 27, 28, 31, 41]. Therefore, Rif1 instead of Rap1 protein is considered in our equations for all the model.

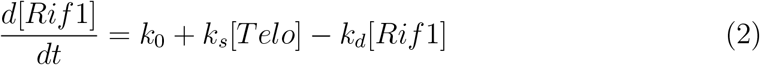

In the equation 2, *k*_0_ is the basal production rate of Rif1, *k*_*s*_ is the synthesis rate induced by activity of telomerase which elongates the telomeres resulting in binding of Rif1 and *k*_*d*_ is the Rif1 degradation rate. Rif1 is telomeric protein concentration and [Telo] is telomerase concentration.We divide the conc. by volume (V) or omega Ω as the number of telomerase is appx 15-20 per cell. Unit for k0 is (nM/min), and for ks and kd is (/min).

We first construct the rate balance plot of the ODE equation 2. Rate balance plot of the ODE equation is a plot of rate of production against the rate of loss. In the ODE equation that is considered rate of synthesis or production, v_1_, which is independent of Telomerase, is given as

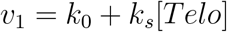

and the degradation term v2 is a function of Rif1 protein.

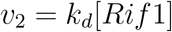

The steady state in the rate balance plot is where v_1_ = v_2_. This steady state or fixed point of [Rif1]_*ss*_ is obtained by taking <u>^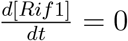^</u>, and its given as

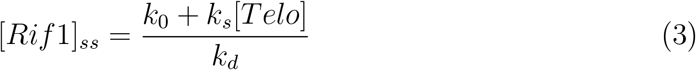

Close form solution for the equation 2 is given by

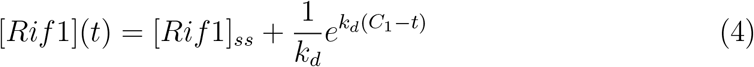

with initial condition *Rif* 1(*t* = 0) = 0 we get 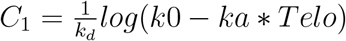

In this 1D model, homeostatic telomere length is obtained by fine tuning the parameters as there are no regulation present in the equation. Due to constant loss and production of nucleotides in telomere, the length of the telomere fluctuates in each cell cycle. To capture this probabilistic variation of telomere length, stochastic simulation is also carried out using Gillespie’s algorithm [15]. Telomere length distribution is calculated for 1000 runs and compared with the deterministic solution.

The simulation results shown in Figure 3(c) shows that the low amount of telomerase cannot not elongates telomeres and telomere length at slightly lower value. TL is maintained between 300-350 bp. High amount of telomerase results in elongation of the telomere length. Refer to Table 1 for parameters of stochastic simulation using Gillespie algorithm [15]and appendix section for more details on stochastic simulation results.

**Figure 1:**
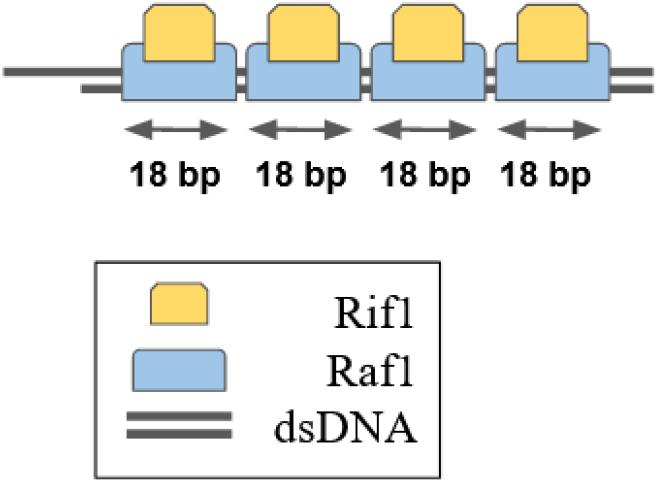
Rif1 protein binds to Rapf1 protein (protein to protein interaction) which further binds to telomeric DNA (protein to DNA interaction).

**Figure 2:**
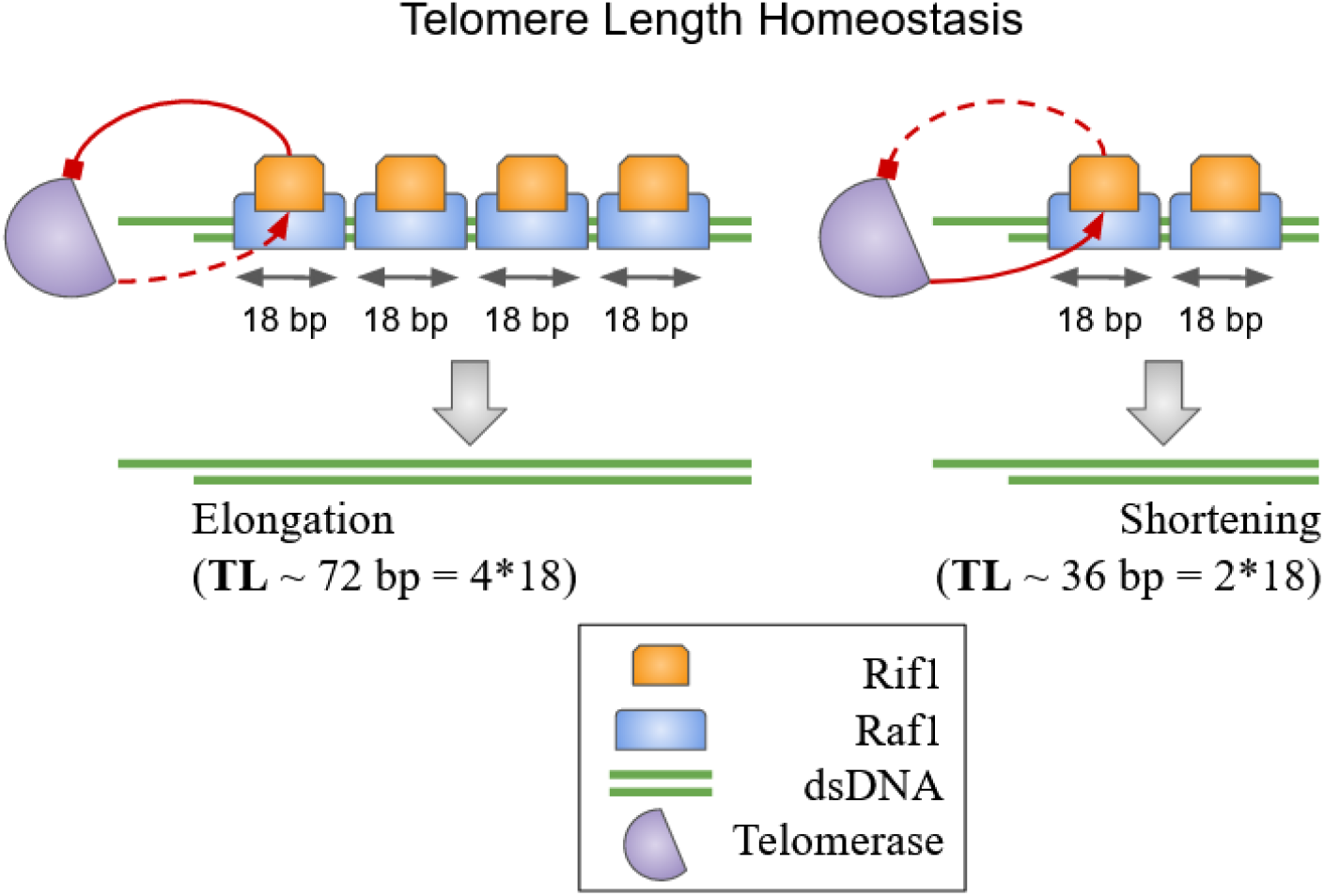
Telomere Length Homeostasis Model diagram. Long telomeres contains more number of Rif1 bind proteins which strongly inhibit telomerase access to the telomeres and gets shorten (inhibition shown with solid red line and activation with dashed red line on the left). Short telomeres has less number of Rif1 bind proteins which weakly inhibit telomerase access to the telomeres and gets elongated (inhibition shown with red dotted line and activation with solid red line on the right).

**Figure 3:**
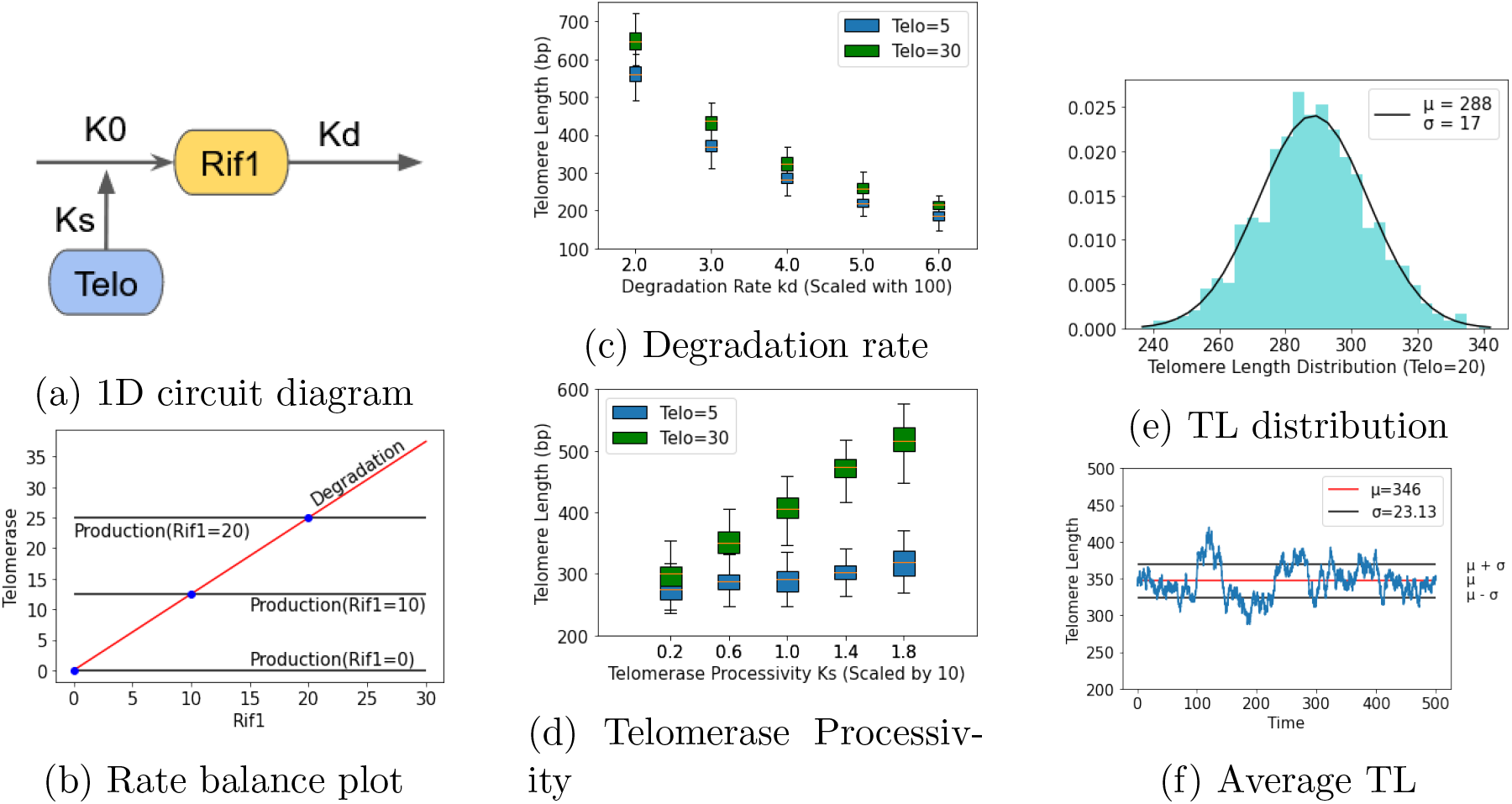
Futile cycle model of telomere regulation by telomerase Circuit diagram with parameters *k*_0_ = 0.8, *k*_*s*_ = 0.06, and *k*_*d*_ = 0.04. (*b*) Rate balance plot between *Rif* 1 and *T elo* plotted for different values of *Rif* 1 = (0, 10, 20). (*c*) Telomere length distribution for different values of degradation rate. For higher degradation rate, mean telomere length is reduced. (*d*) Telomere length distribution for different values of telomerase processivity. For higher telomerase telomere length distribution shifts to higher value compared to slight change for lower value. The (*e*) histogram plot of telomere length distribution (TL) obtained over 1000 iteration of stochastic simulation for Telo=0 (no telomerase) with mean TL *µ* = 288 bp and *σ*= 17 using TL equation 1 for Ω = 20. (*f*) Deterministic and stochastic simulation starting with initial condition Rif1(t=0)=0 reaching the steady state expression given by 3.

**Table 1:**
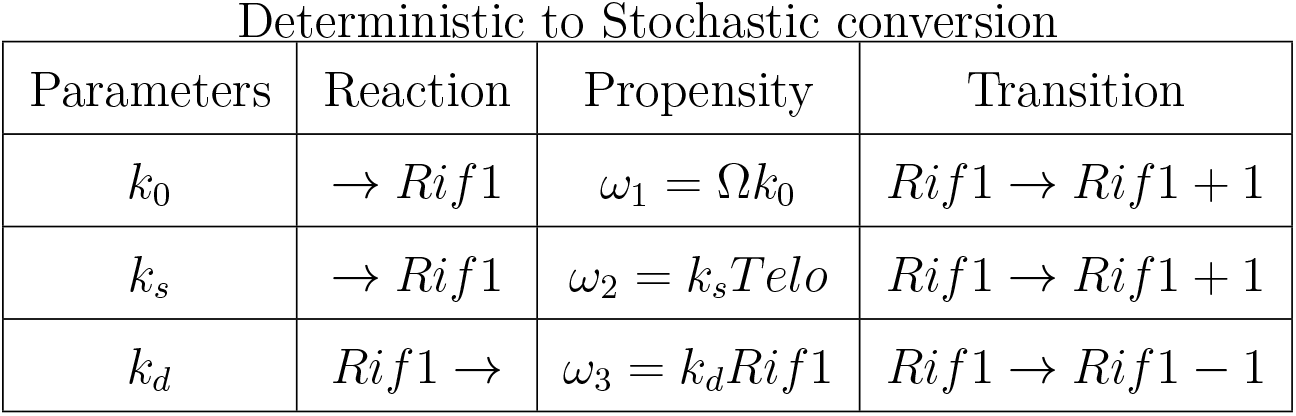
Stochastic version of 1D Model

| Parameters | Reaction | Propensity | Transition |
| --- | --- | --- | --- |
| $k_0$ | $\rightarrow Rif1$ | $\omega_1 = \Omega k_0$ | $Rif1 \rightarrow Rif1 + 1$ |
| $k_s$ | $\rightarrow Rif1$ | $\omega_2 = k_s Telo$ | $Rif1 \rightarrow Rif1 + 1$ |
| $k_d$ | $Rif1 \rightarrow$ | $\omega_3 = k_d Rif1$ | $Rif1 \rightarrow Rif1 - 1$ |

Significance of 1D model is to provide a broad idea about the variation of telomere length distribution in somatic cells in the presence of low to high telomerase concentration. Telomerase can add nucleotides up to hundred base pairs in *S*.*cerevisiae* [55], and it varies from each cycle of cell division. Therefore telomerase processivity has impact on homeostasis length. Lowering the telomerase concentration leads to a faster decrease in telomere length. In somatic cells telomerase concentration appears to be very low. Normal human somatic cells have telomerase levels below the level required for telomere maintenance and their telomeres shorten with each cell division [40]. Telomere length below a critical length can trigger a p53 dependent checkpoint response leading to cell cycle arrest [8, 9, 25]. Based on the evidence [8] that ability of cells to proliferate is limited by short telomeres and gradual shortening in somatic cells where low telomerase concentration leads to finite proliferation capacity [12].

1D model is a phenomenological model that relates proliferative potential of a cell to the amount of telomerase present in the cell. This is shown mathematically by [6]. Blagoev showed through a mathematical model that proliferative capacity of a cell grows exponentially with the telomerase concentration. There exists two types of telomerase-telomere interaction. First is when telomerase concentration is higher, many telomerase complexes are available for elongation of each telomere and whenever telomere is shorter, telomerase is available in sufficient quantity to associate and elongate each telomere. Second is when telomerase concentration is lower than the number of short telomeres, an adaptive control mechanism exists in which telomere length is limited by the amount of telomerase. Telomere length homeostasis depends on the proximity of telomerase near telomere.

Its a simple model which explains the TL distribution based on fine tuning the parameters. There is no regulation involved in 1D model. 1D model did not provide enough insights on regulatory behavior of Rif1 on Telomerase or vice versa. It does not explain the dynamics of elongation and shortening of telomeres. As there is no regulation or feedback mechanism involved, parameters are not robust for a wide range of values in the 1D model.

### A 2D model with a negative feedback loop

Though 1D linear model provided a basic idea and mathematical structure about the teleomeric homeostatic length mechanism, it is dependent on kinetic parameters. Kinetic parameters are subjected to variations and therefore, cannot play a strong role in maintaining the length homeostatsis. We expanded the 1D-model by taking telomerase as a variable rather than constant and involves in regulating telomere bound protein Rif1 through a negative feedback loop (Figure 4a). This is based on the information from [37] that there is an inverse relationship exists between the telomerase activity and telomere length for the budding yeast. It is also known from the protein counting model (PCM) [39] that a high amount of Rif1 protein inhibits telomerase access to telomeres whereas for a very low concentration of Rif1 protein, inhibition tends to be mild. Based on this information about the negative regulation of Rif1 protein by telomerase (Telo) we construct a two-dimension model with a negative feedback as follows.

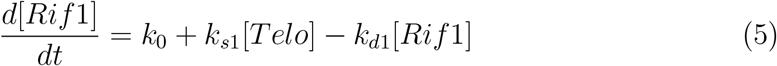

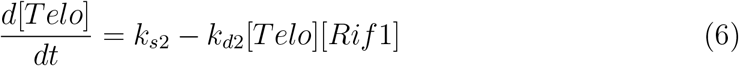

**Figure 4:**
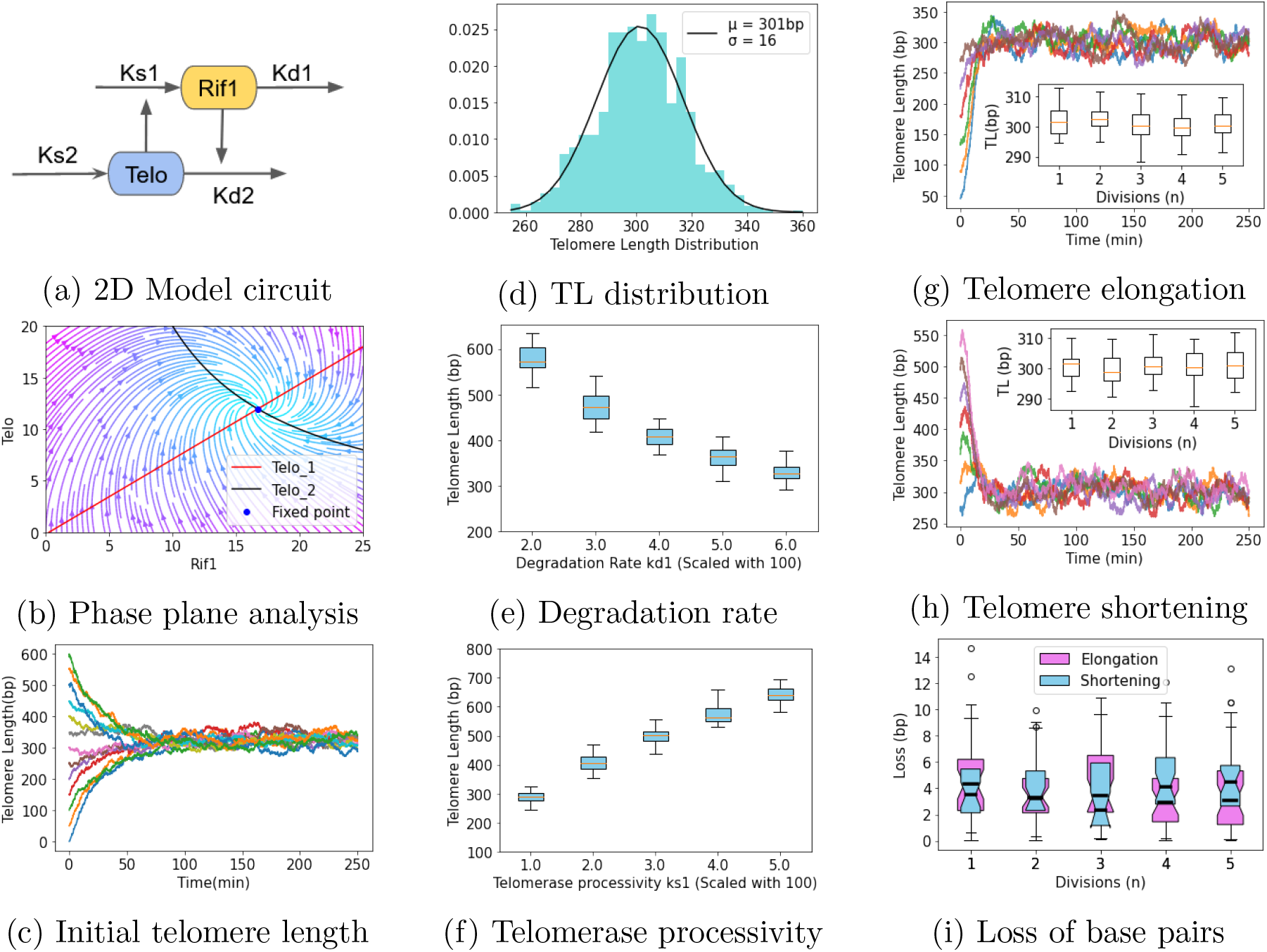
2D model of regulation by Rif1 and Telomerase. (*a*) Circuit diagram. Parameters chosen for the model are *k*_0_ = 0.02, *k*_*s*1_ = 0.11, *k*_*d*1_ = 0.08, *k*_*s*2_ = 2, *k*_*d*2_ = 0.01. (*b*) Phase plane analysis of the model. Null clines *T elo*_1_ and *T elo*_2_ mentioned in equation 7 intersects at a stable fixed point shown by blue. (*c*) Stochastic simulation trajectories of the 2D model describes the behavior of telomere at each cell division with different initial telomere length at (t=0). Parameters for the stochastic simulations are given in Table 2. Short telomeres get elongated while long telomeres are shortened and reaches steady stable length of*≈* 300bp over few cell divisions. (*d*) Histogram plot of telomere length distribution with mean *µ* = 301bp and std deviation *σ*= 16 obtained by fitting normal distribution function over TL histogram. (*e*) Impact of variation in kinetic parameter *k*_*d*1_ as degradation rate on telomere length distribution. (*f*) Impact of variation in kinetic parameter *k*_*s*1_ as telomerase processivity on telomere length distribution. (*g*) Elongation of telomeres over 5 cell divisions. (*h*) Shortening of telomeres over 5 cell divisions. (*i*) Loss of base pairs during elongation and shortening is within *≈* 3bps over few cell divisions.

**Table 2:** Stochastic version of 2D Model

| Parameters | Reaction | Propensity | Transition |
| --- | --- | --- | --- |
| $k_0$ | $\rightarrow Rif1$ | $\omega_1 = \Omega k_0$ | $Rif1 \rightarrow Rif1 + 1$ |
| $k_{s1}$ | $\rightarrow Rif1$ | $\omega_2 = k_{s1}Telo$ | $Rif1 \rightarrow Rif1 + 1$ |
| $k_{d1}$ | $Rif1 \rightarrow$ | $\omega_3 = k_{d1}Rif1$ | $Rif1 \rightarrow Rif1 - 1$ |
| $k_{s2}$ | $\rightarrow Telo$ | $\omega_4 = \Omega k_{s2}$ | $Telo \rightarrow Telo + 1$ |
| $k_{d2}$ | $Telo \rightarrow$ | $\omega_5 = (1/\Omega)k_{d2}TeloRif1$ | $Telo \rightarrow Telo - 1$ |

In the above equation 5, *k*_0_ is the basal Rif1 production rate, k_*s*1_ is the rate of synthesis induced by telomerase which elongates the telomeres that results in binding of Rif1 over few cell divisions, and *k*_*d*1_ is the Rif1 degradation rate. Further in equation 6, *k*_*s*2_ is rate of telomerase production and *k*_*d*2_ is degradation of telomerase carried out by Rif1 protein. The 2D model consists of positive regulation of Rif1 by telomerase and negative regulation of telomerase by Rif1 that generates a negative feedback loop. Negative feedback loops are known to provide rapid response and homeostasis [1]. Equations 5 and 6 captures the molecular dynamics quite well compared to equation and (4) provided in the supplementary material of [62]. It is able to generate and fit the experimental data of telomere length distribution [55] quite well.

Since the 2D-ODE is a coupled nonlinear ODE, we performed nullcline analysis by taking *d*[*Rif* 1]*/dt*=0 and = *d*[*T elo*]*/dt*=0. The algebraic equations are

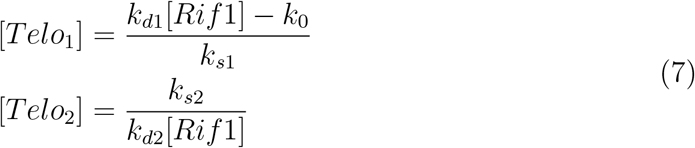

[Telo_1_] and [Telo_2_] are the algebraic equations of 2D-ODE and the nullclines of these two equations are shown in Figure 5b. For the choice parameters, [Rif1] and [Telo] meets at *≈* (16.7, 12). It can also be seen that due to the presence of negative feedback loop the variables settles down to the steady state with a damped oscillations. No damped oscillations exists here, initially when Rif1 is low, Telo increases but when Rif1 reaches a particular level sufficient enough, inhibits telomerase activity and settles to a steady state.

**Figure 5:**
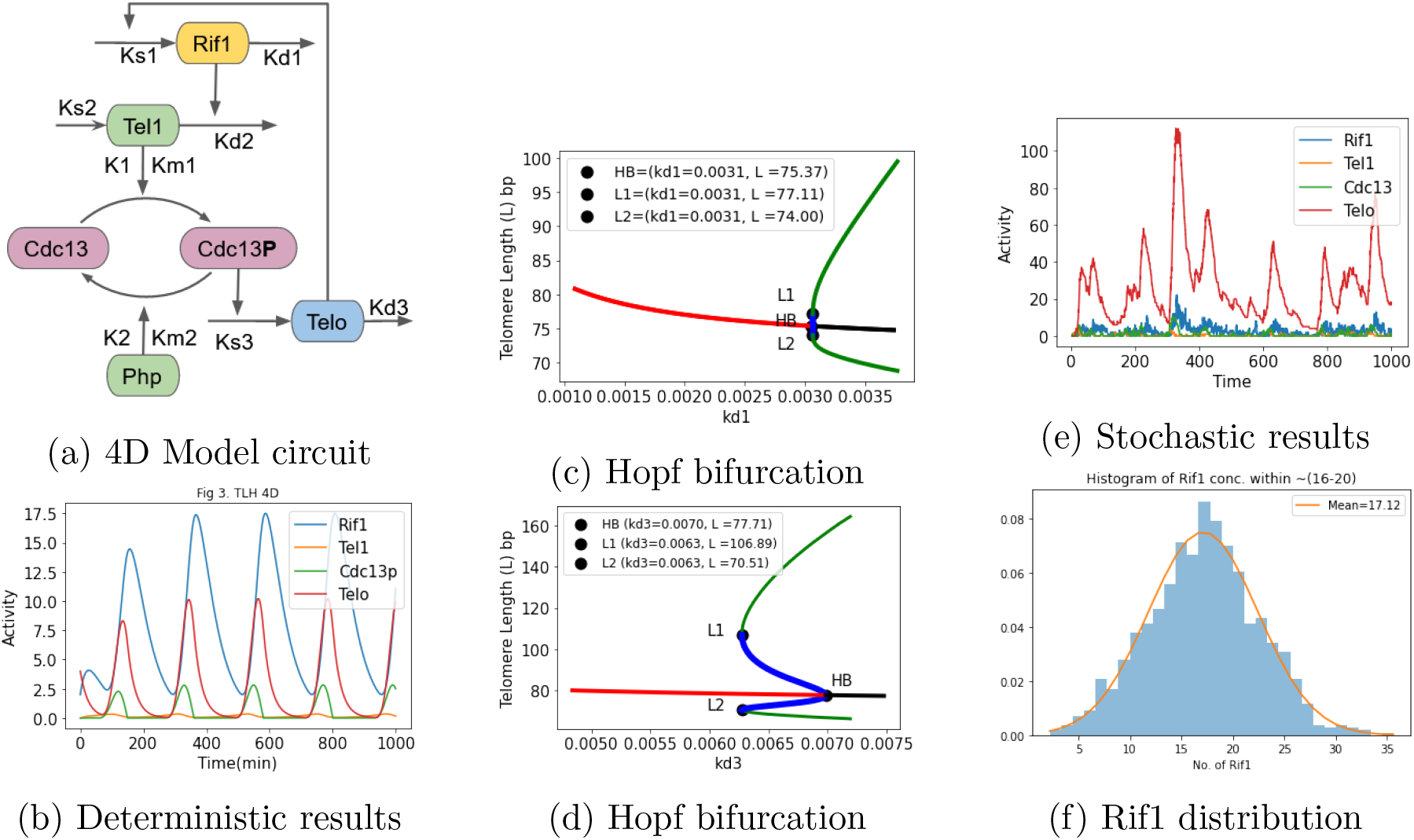
4D Model with sustained oscillations (*a*) Circuit diagram of the model with addition of Cdc13 regulated by Tel1 kinase. Parameter for the model are *k*_0_ = 0.02, *k*_*s*1_ = 0.06, *k*_*d*1_ = 0.02, *k*_*s*2_ = 0.01, *k*_*d*2_ = 0.01, *k*_*s*3_ = 0.2, *k*_*d*3_ = 0.04, *k*_1_ = 0.8, *k*_2_ = 0.02, *Km*_1_ = 0.002, *Km*_2_ = 0.002, *n* = 1, *Php* = 10, *Cdc*13_*T*_ = 10 with initial condition Rif1(t=0)=3, Tel1(t=0)=0, Cdc13P(t=0)=0, Telo(t=0)=4. (*b*) Addition of Cdc13p activation by Tel1 kinase added a delay acting as intermediate species resulting in proper oscillations. Simulation results in oscillatory response of the circuit elements indicating Rif1 oscillating between higher and lower values which indirectly relates oscillation in telomere length.(*c*) Supercritical Hopf bifurcation. One parameter bifurcation diagram plot for the parameter *k*_*d*1_ = 0.003 and telomere length (TL). (*d*) Subcritical Hopf bifurcation. One parameter bifurcation diagram plot for the parameter *k*_*d*3_ = 0.006 and telomere length (TL). (*e*) Stochastic simulation results obtained using parameters given in the Table 3 and using equation 1. (*f*) Histogram plot of bound Rif1 proteins obtained from stochastic simulation with a mean *µ* = 17 no. of Rif1 proteins.

We also performed stochastic simulations of the 2D model to determine the mean length and its variations. In S.*cerevisiae* basal telomerase loss is 3 nts per generation [37]. We carried out for low and high Rif1 initial protein concentration (correspondingly low and high telomere length) to determine the time taken for the 2D model reach the mean-telomeric length. The mean telomeric-length after 1000 number of iterations is 301 ± 16 bp. This mean-length is very close to the experimental value for S.*cervisiae* as shown by Xu et al.[62]. Similar results have also been obtained by [6] using the Hill’s function to determine the mean-telomeric length. However, our model is different from their model from the way we have fully simulated based on the mechanistic details.

**Table 3:**
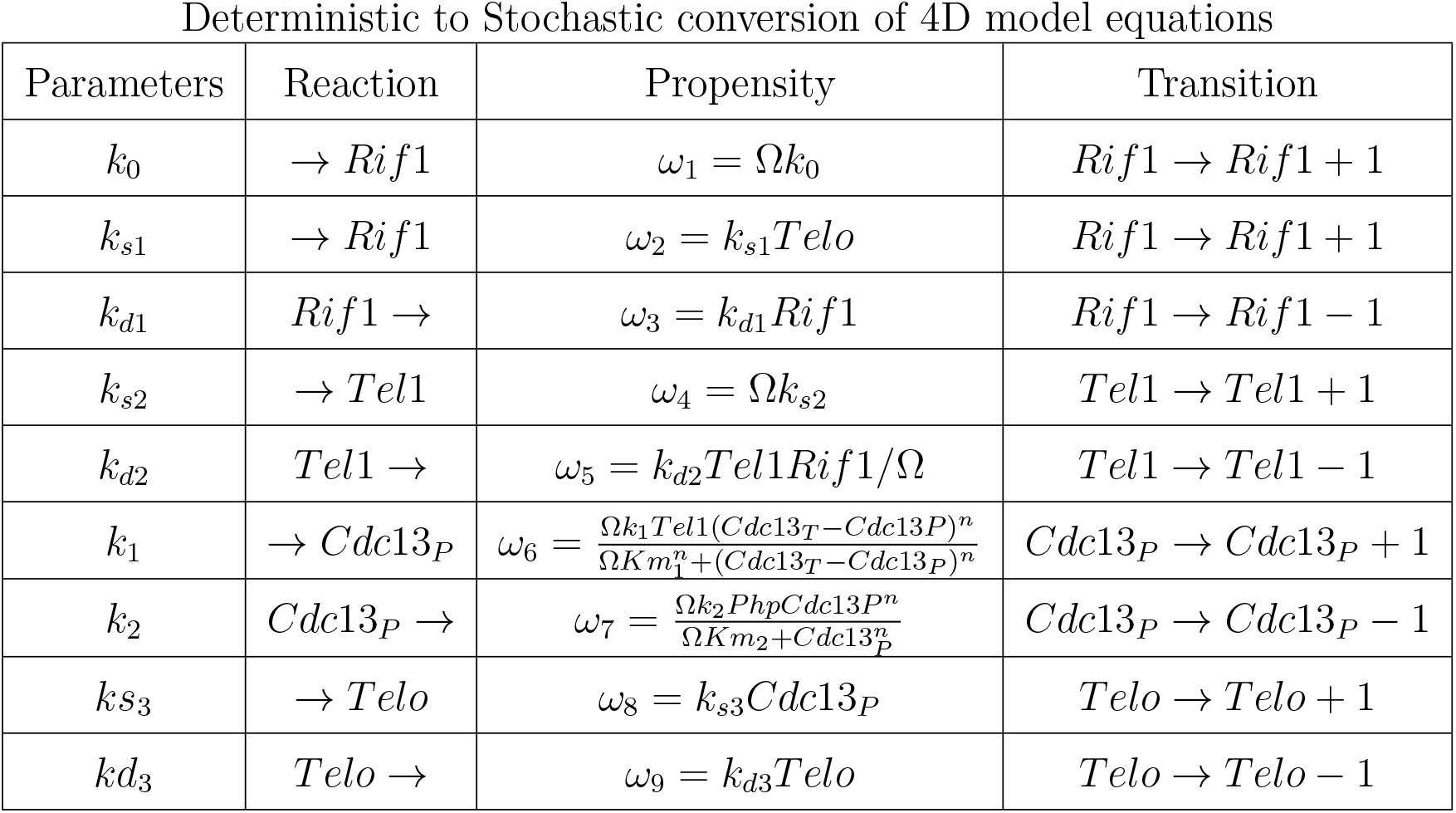
Stochastic version of 4D Model

| Parameters | Reaction | Propensity | Transition |
| --- | --- | --- | --- |
| $k_0$ | $\rightarrow Rif1$ | $\omega_1 = \Omega k_0$ | $Rif1 \rightarrow Rif1 + 1$ |
| $k_{s1}$ | $\rightarrow Rif1$ | $\omega_2 = k_{s1}Telo$ | $Rif1 \rightarrow Rif1 + 1$ |
| $k_{d1}$ | $Rif1 \rightarrow$ | $\omega_3 = k_{d1}Rif1$ | $Rif1 \rightarrow Rif1 - 1$ |
| $k_{s2}$ | $\rightarrow Tel1$ | $\omega_4 = \Omega k_{s2}$ | $Tel1 \rightarrow Tel1 + 1$ |
| $k_{d2}$ | $Tel1 \rightarrow$ | $\omega_5 = k_{d2}Tel1Rif1/\Omega$ | $Tel1 \rightarrow Tel1 - 1$ |
| $k_1$ | $\rightarrow Cdc13_P$ | $\omega_6 = \frac{\Omega k_1 Tel1 (Cdc13_T - Cdc13_P)^n}{\Omega K m_1^n + (Cdc13_T - Cdc13_P)^n}$ | $Cdc13_P \rightarrow Cdc13_P + 1$ |
| $k_2$ | $Cdc13_P \rightarrow$ | $\omega_7 = \frac{\Omega k_2 Php Cdc13_P^n}{\Omega K m_2 + Cdc13_P^n}$ | $Cdc13_P \rightarrow Cdc13_P - 1$ |
| $k_{s3}$ | $\rightarrow Telo$ | $\omega_8 = k_{s3}Cdc13_P$ | $Telo \rightarrow Telo + 1$ |
| $k_{d3}$ | $Telo \rightarrow$ | $\omega_9 = k_{d3}Telo$ | $Telo \rightarrow Telo - 1$ |

We have also attributed our kinetic parameters *k*_*s*1_ to processivity and *k*_*d*1_ to telomeric loss during cell division. The parameter robustly captures these aspects. For example, figure 4(c) shows that the mean telomere length depends on the degradation rate of telomere bound proteins Rif1. If Rif1 protein gets degraded quickly due to external or internal stress factors then sufficient amount of Rif1 is not available for it to bind to telomeres for protection.

It also cause quick loss of telomeric DNA and results in a shift of mean telomeric length to a lower value. It also suggests about the possibility that Rif1 pathway may be getting exploited by stem cells or cancer cells to keep high telomere length. Similarly figure 4(d) indicates that the high amount of telomerase in cells, particularly near the telomeric end results in the shift of the mean telomere length to a higher value. We compare this results with that of [62] (refer to supplementary section of figures 2B and 2C of [62] for more details).

### 4D Model with Telomere Length Oscillations

We extended the 2D model by adding the Tel1 kinase, a third dynamical variable giving rise to a 3D model. Since this gave a very similar result as that of 2D, the details are relegated to supplementary material. We extended this 3D model by adding Cdc13, a single stranded telomeric DNA binding protein that positively regulates yeast telomere replication by recruiting telomerase [10]. Association between Est1 telomerase subunit and Tel1 kinase mediated phosphorylation of the Cdc13 telomerase recruitment domain is essential for the recruitment of telomerase. Mathematical modeling of this underlying molecular mechanism captures the essential interactions among these telomeric species Rif1-Tel1-Cdc13-Telo. Telomerase recruitment by Cdc13 creates a Time delay. System is is given by following equations:

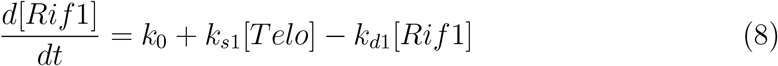

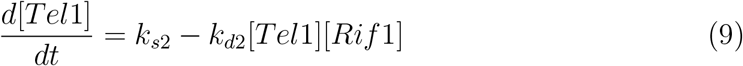

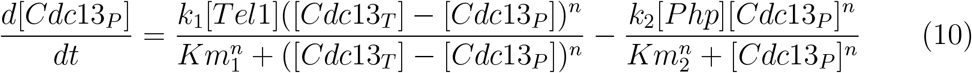

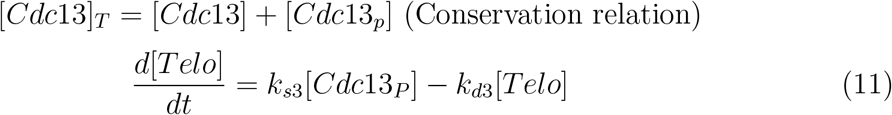

In equation 8, *k*_0_ is the basal Rif1 production rate, *k*_*s*1_ is the synthesis rate induced by activity of telomerase which elongates the telomeres resulting in binding of Rif1 over few cell divisions, and *k*_*d*1_ is Rif1 degradation rate. In equation 9, *k*_*s*2_ is Tel1 kinase production rate and *k*_*d*2_ is Rif1 mediated Tel1 kinase degradation rate. In equation 10, *k*_1_ is Cdc13 phosphorylation rate mediated by Tel1 kinase, *k*_2_ is Cdc13 dephosphorylation rate mediated by Php phosphatase, *Cdc*13_*T*_ is total amount of Cdc13, *Cdc*13_*P*_ is phohphorylated Cdc13, *Km*_1_ and *Km*_2_ are Michaelis–Menten rate constants,’n’ is the Hill’s coefficient. In equation 11, *k*_*s*3_ is phosphorylated Cdc13 mediated telomerase production rate and *k*_*d*3_ is telomerase degradation rate.

### Bifurcation and stochastic simulations

Deterministic mathematical simulations provide an initial insight into understanding the behavior of the system by modeling it. However, in the present case, the gain and loss of telomere is highly variable in each cell divisions, we performed stochastic simulations. As a first step, we converted the deterministic kinetic constants in the 4D model into a stochastic constants and performed simulations using Gillespie algorithm [16]. We are interested particularly to generate the distribution of Rif1 concentration that is later shown to relate to the telomere length. Sufficient delay was introduced by the addition of Tel1 kinase mediated phosphorylation of Cdc13 between negative regulation of telomerase by Rif1. This negative feedback with delay results in giving rise to sustained limit cycle oscillation of Rif1 and Telo as shown in figure 5b In figure 5c when *kd*_1_ *<* 0.003, systems remains at a steady state and behaves like a stable spiral but beyond *kd*_1_ *>* 0.003 the system bifurcates and converts into a limit cycle. From the bifurcation simulation, there exists a critical telomere length (L) 72-108 bp which is equivalent to 4-6 no. of Rif1 which which must remain bounded to telomeric DNA to maintain telomere length homeostasis. Telomere length (L) below a critical length, leads to activation of DNA damage-checkpoint pathways and cellular senescence or apoptosis [7]. Our simulation results matches with the experimentally determined length of the shortest telomere TL 90bp from [55]. Based on the earlier findings of that telomere length switches between an extendible and non extendible states, we found out that the telomeres length oscillates between elongation and shortening states based on the molecular mechanisms.

## Conclusions

This research provides a mathematical model for the mechanisms that regulate telomere length at the molecular level for budding yeast *S. cerevisiae*. We have designed a biological circuit step by step to describe telomere length homeostasis and developed simple ODE models of one dimension to four dimensions. Then we analyzed the models using dynamical systems theory. To get a realistic picture stochastic simulations were done using Gillespie algorithms to relate the number of telomeres bound proteins and telomerase with previous experimental understandings and explain the oscillation of telomere length from elongation to shortening state during cell division. [62] has shown in their simulation based on geometric distribution model similar mean-telomeric length. They started with chemical kinetic equations and converted it to a geometric probabilistic model, given in supplementary [62]. We have modeled on the same lines as shown by them except that we have included negative feedback loop in the model. Though our model is phenomenological to a large extent, it is amenable to analysis and provides insight about the loss of telomeric length and processivity in terms of kinetic parameters. Telomere length homeostasis mechanism across different model species is conserved. It will be interesting to develop and extend models for other organisms. Next, we would like to see the role of DNA replication in telomere length maintenance and build upon the hypothesis proposed by [20] to develop a replication fork based mathematical model in the future.

## Future Work

Recent studies for the yeast *S. cerevisiae* have discovered that telomerase can be selectively turned on and off [56] referring to assembly and disassembly of the subunits of telomerase. Initially, the telomerase complex is present in a poised state with a missing subunit and during the telomeric DNA replication missing subunit attaches to form a fully active telomerase complex. The assembly pathway, which is driven by the interaction of the Est3 telomerase subunit with a previously formed Est1–TLC1–Est2 preassembly complex, is highly regulated, involving Est3-binding sites on both Est2 and Est1 as well as an interface on Est3 itself that functions as a toggle switch. Cdc13 phosphorylation of the Est1 subunit helps in mediating the telomerase access to telomeres. Cancer cells might be using this switch of Est1 phosphorylation or dephosphorylation to activate the complex to keep telomerase levels high to maintain longer telomeres. Cancer cells may be regulating this circuit. We further need to investigate the activation of the P53 pathway by short telomeres. This model can be extended further to explain mammalian telomere length regulation in future.

## Acknowledgements

This work was done as a part of masters thesis research work supported by the Center of Computational Biology, IIIT Delhi, India.

