## Supplementary Material for "Mechanistic Model of Telomere Length Homeostasis in Yeast"

### Mechanistic Model of Telomere Length Homeostasis in Yeast Supplementary Material

Ghanendra Singh[1] and Sriram K.[1]

#### Extension of 2D to 3D Model: Role of Tel1 kinase

Though 2D model with negative feedback loop explained the telomere length homeostasis, we decided to unpack the mechanism to introduce other players responsible for the dynamics. Smogorzewska et al [3] have shown that the two DNA damage response kinases Tel1 and Mec1 [1] may be responsible for maintaining the homeostatic length through regulatory interactions with telomere and telomerase.

Role of negative feedback loop was explained by 2D model though it does not explain about how it exists between telomerase and telomeres. Telomere length regulation by telomeric proteins and the possible role of DNA damage response kinases Tel1 and Mec1 [1] for telomere length control was mentioned by [3]. In this 3D model we have considered the role of Tel1 kinase which recruits telomerase to telomeres. A simple three-dimensional system of equations is given by:

$$\frac{d[Rif1]}{dt} = k_0 + k_{s1}[Telo] - k_{d1}[Rif1] \tag{1}$$

$$\frac{d[Tel1]}{dt} = k_{s2} - k_{d2}[Tel1][Rif1] \tag{2}$$

$$\frac{d[Telo]}{dt} = k_{s3}[Tel1] - k_{d3}[Telo] \tag{3}$$

In equation 1, $k_0$ is the basal Rif1 production rate, $k_{s1}$ is the synthesis rate induced by activity of telomerase which elongates the telomeres resulting in binding of Rif1 over few cell divisions, and $k_{d1}$ is Rif1 degradation rate. Further in equation 2, $k_{s2}$ is Tel1 kinase production rate and $k_{d2}$ is Rif1 mediated Tel1 kinase degradation rate. In equation 3, $k_{s3}$ is Tel1 kinase associated telomerase production rate and $k_{d3}$ is telomerase degradation rate. 3D model is obtained by addition of Tel1 kinase which is negatively regulated by Rif1. Rif1 recruits PP1 to inhibit Tel1 kinase [2]. Simulation results shows that the circuit produces initially damped oscillation and then quickly reaches a steady-state as shown in figure 1b. Tel1 kinase dynamics are faster. Possibly the delay comes from phosphorylation of Cdc13 by Tel1 to recruit telomerase. Model gives insights about existence of an oscillator if there exist sufficient delay with negative feedback.

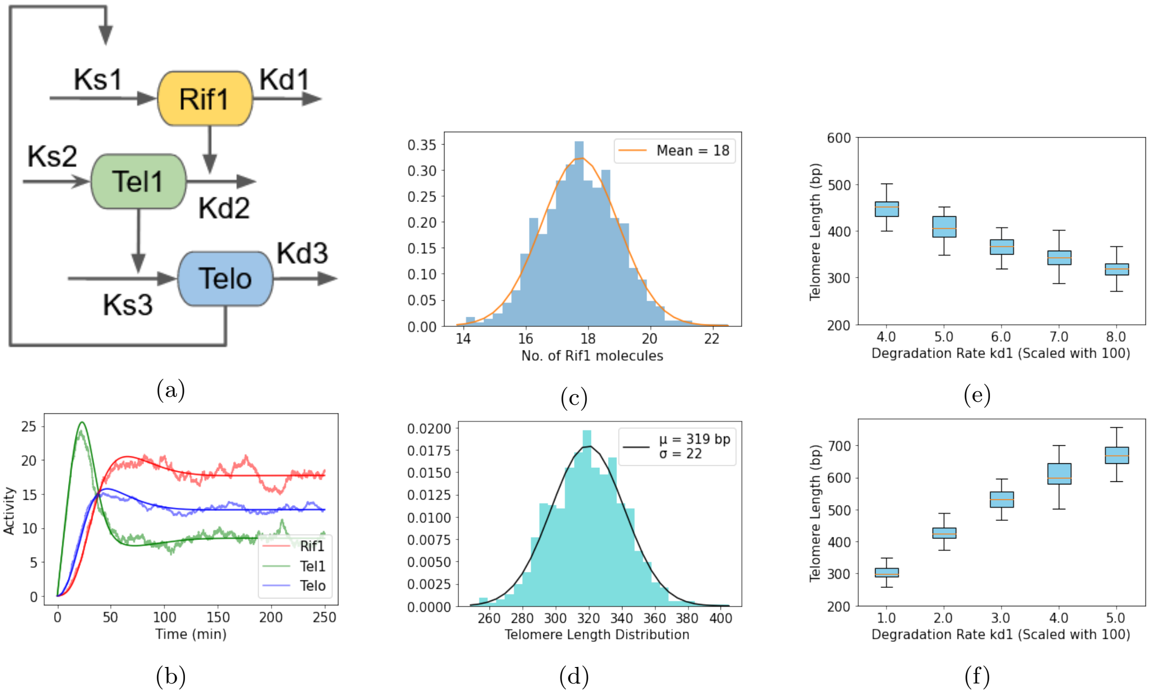


Figure 1: 3D Model with damped oscillations
($a$) Circuit diagram with inclusion of Tel1 kinase. Parameters for the 3D model are $k_0 = 0.02, k_{s1} = 0.1, k_{s2} = 1.5, k_{d2} = 0.01, k_{s3} = 0.03, k_{d3} = 0.02$ with initial values Rif1(t=0)=0,Tel1(t=0)=0,Telo(t=0)=0. ($b$) Deterministic and stochastic simulation results initially in damped oscillations reaching to steady state. ($c$) Histogram plot of no. of Rif1 protein with mean $\mu = 18$. ($d$) Histogram plot of telomere length distribution (TL) with $\mu = 319$bp and $\sigma = 22$ obtained over 1000 iteration of stochastic simulation for $\Omega = 20$. ($e$) Impact of variation in kinetic parameter $k_{d1}$ as degradation rate on telomere length distribution. For higher value of degradation rate, mean telomere length is reduced. ($f$) Impact of variation in kinetic parameter $k_{s1}$ as telomerase processivity on telomere length distribution. For high telomerase processivity mean telomere length distribution shifts to a higher value. Median is indicated by the horizontal line in boxes.
